# Visualization of trigeminal ganglion sensory neuronal signaling regulated by Cdk5

**DOI:** 10.1101/2021.05.07.443189

**Authors:** Minghan Hu, Andrew Doyle, Kenneth M. Yamada, Ashok B. Kulkarni

**Affiliations:** Functional Genomics Section and Cell Biology Section, National Institute of Dental and Craniofacial Research, National Institutes of Health, Bethesda, MD, USA.; Cell Biology Section, National Institute of Dental and Craniofacial Research, National Institutes of Health, Bethesda, MD, USA.; Cell Biology Section, National Institute of Dental and Craniofacial Research, National Institutes of Health, Bethesda, MD, USA..gov; Functional Genomics Section, National Institute of Dental and Craniofacial Research, National Institutes of Health, Bethesda, MD, USA.

**Keywords:** Allodynia, Cdk5, GCAMP6f, inflammation, in vivo imaging, neuron, nociceptor, pain, trigeminal ganglia, TRPV1

## Abstract

The mechanisms underlying facial and oral pain are still incompletely understood, posing major therapeutic challenges. Cyclin-dependent kinase 5 (Cdk5) is a key neuronal kinase involved in pain signaling. However, the regulatory roles of Cdk5 in orofacial pain signaling and the possibility of therapeutic intervention at the level of mouse trigeminal ganglion primary neurons remain elusive. In this study, we used optimized intravital imaging to directly compare trigeminal neuronal activities after mechanical, thermal, and chemical stimulation. We then tested whether facial inflammatory pain in mice could be alleviated by the Cdk5 inhibitor peptide TFP5. We demonstrated regulation of total Ca^2+^ intensities by Cdk5 activity using transgenic and knockout mouse models. In mice with orofacial inflammation, application of TFP5 specifically decreased total Ca^2+^ intensities in response to noxious stimuli. It also alleviated inflammation-induced allodynia by inhibiting activation of trigeminal peripheral sensory neurons. Cdk5 inhibitors may provide promising non-opioid candidates for pain treatment.

## Introduction

Pain affecting the face or oral cavity (orofacial pain) afflicts 5%-12% of the population worldwide and can severely decrease the quality of life (Crandall, 2018; Herrero Babiloni et al., 2018). It also contributes to other health and social problems such as the opioid crisis (Sugawara et al., 2019). Clinicians struggle to alleviate pain due to our current inadequate understanding of the detailed mechanisms. Understanding trigeminal pain signal coding and transmission will be paramount for the effective management of orofacial pain.

The trigeminal ganglion (TG) of the peripheral nervous system mediates orofacial pain signaling. It contains nociceptor cell bodies that innervate and contribute to pain sensing within the face and oral cavity (Kim et al., 2014; Rothermel et al., 2011), transmitting pain sensation through the TG to the central nervous system (CNS) (Bista and Imlach, 2019). Both noxious and non-noxious stimuli can activate specific nociceptors in TG neurons, such as thermal, mechanical, and chemical receptors. Among the nociceptors of the TG, the transient receptor potential vanilloid type-1 (TRPV1) plays a key role in pain perception. TRPV1 is a member of the TRP channel family and is a calcium-permeable channel. It is primarily expressed in small- to medium-sized nociceptor neurons of the TG and dorsal root ganglia. TRPV1 is sensitive to various stimuli, including thermal, chemical, and mechanical stimuli (Gonzalez-Ramirez et al., 2017; Julius, 2013; Warwick et al., 2019). The activity of TRPV1 can be regulated by phosphorylation, which results in increased sensitivity to pain stimuli (Hall et al., 2018; Jendryke et al., 2016).

Recently, increasing interest has focused on the role of cyclin-dependent kinase-5 (Cdk5) in regulating pain signaling. Cdk5 is a proline-direct serine/threonine kinase that is expressed ubiquitously (Hellmich et al., 1992; Liu et al., 2016). When bound to its activators p35 or p39, Cdk5 phosphorylates target proteins that participate in maintaining normal neuronal development and homeostasis (Cortes et al., 2019; Pareek et al., 2013). Cdk5 is also a key component of the nociceptive pathway, where its expression and activity is increased upon inflammation following nociceptive stimulation (Pareek et al., 2006; Yang et al., 2007). Intrathecal administration of an inhibitor of several Cdks can attenuate both formalin-induced nociceptive responses and diminish morphine tolerance (Wang et al., 2005; Wang et al., 2004). Previously, our laboratory validated the *in vivo* roles for Cdk5 in orofacial hypoalgesic or hyperalgesic behavior in transgenic mice with decreased or increased Cdk5 activity, respectively (Prochazkova et al., 2013).

Cdk5 is known to phosphorylate TRPV1 at Thr-407 (mouse or human) (Liu et al., 2015) or Thr-406 (rat) (Pareek et al., 2007) to enhance channel function. Because Cdk5 is associated with pathological events in inflammation and pain, pharmacological targeting of this kinase provides an attractive prospect for potential pain therapy. However, the roles of Cdk5 in the cellular basis of sensory pain coding remain elusive, e.g., in specific (unimodal) or polymodal neuronal signaling. In addition, the activities of peripheral nociceptors are often studied using sensory nerve recordings in *ex vivo* tissue preparations, and technical barriers have limited in vivo investigation of the role of primary nociceptors and neuronal pain coding in sensitization and hyperalgesia. For example, facial pain in response to light touch (allodynia) needs more mechanistic characterization to define the neuronal signaling, e.g., in terms of specific/unimodal or polymodal signaling and its magnitude.

In this study, we applied intravital calcium imaging to characterize the response patterns to multiple different stimuli of individual and populations of neuronal activities in the TG of living mice. We labeled TRPV1-linage neurons with GCamp6f to simultaneously monitor their responses to sequential non-noxious and noxious stimuli applied to the orofacial area of mice with different genetic backgrounds or after pharmacological treatment.

One goal of our study was to characterize the nature of TG signaling in comparisons of non-painful and painful stimuli in terms of the specificity or overlap of signaling to different modes of stimuli. Another goal was to identify the effects of levels of Cdk5 activity on this orofacial somatic sensing and pain perception for different types of stimuli by examining alterations in calcium signaling and involvement of individual neurons. Additionally, we tested whether a specific Cdk5 inhibitor could alleviate inflammation-induced allodynia or hyperalgesia signaling from primary nociceptors. We discovered that enhanced Cdk5 activity in TG not only increased calcium influx in individual neurons, but also expanded the population size of nociceptive cells that responded to orofacial pain. In contrast, genetically decreased Cdk5 activity in mice resulted in hyposensitivity. Notably, Cdk5 inhibitor application either locally to the TG or by systemic administration attenuated inflammation-induced nociceptor hyperactivity in the TG. These findings may have future translational impact by identifying a peripheral, rather than a CNS, therapeutic target for orofacial pain.

## Results

### In vivo imaging of trigeminal ganglion nociceptor responses to orofacial mechanical, thermal, or chemical pain

We visualized trigeminal pain signaling directly using TRPV1-cre mice (Cavanaugh et al., 2011) crossed with Rosa26-CAG-flox-stop-GCaMP6f mice to genetically restrict expression of the calcium indicator GCaMP6f to TRPV1-linage neurons. For monitoring dynamic neuronal activities of the trigeminal ganglion (TG) in real time as it responds to orofacial stimulation and pain, we adapted an intravital in vivo live imaging technique (Ghitani et al., 2017) to expose the TG and optically record activity from large ensembles of genetically encoded primary sensory neurons (Fig.1 A-B).

**Fig. 1.**
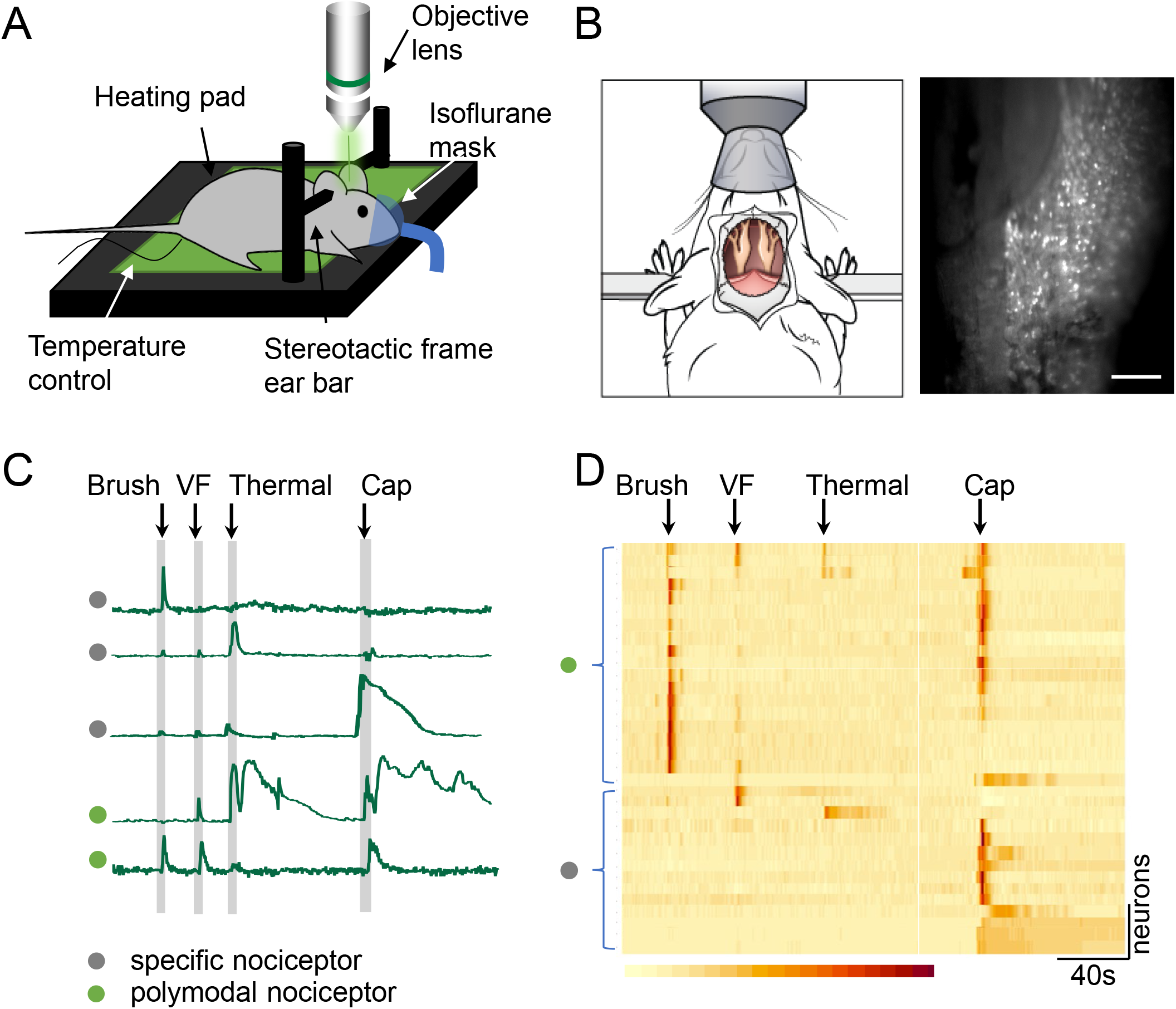
In vivo imaging of trigeminal ganglia in response to different modes of orofacial pain stimulation. A. Diagram of the surgical mount for a mouse with imaging system in preparation to expose the trigeminal ganglion. B. A top-down view of the mouse surgical preparation with exposed trigeminal ganglia imaged at 4X magnification (left), demonstrating visualization of responses induced by pinch stimulation (right). Scale bar, 500 μm. C. Five representative Ca^2+^ intensity traces for different types of nociceptors observed during live imaging. The first three traces represent specific nociceptor responses to brush, thermal (47°C), or capsaicin stimuli, respectively. The bottom two Ca^2+^ traces illustrate polymodal responses. VF, von Frey hair fiber. Cap, capsaicin. D. Heatmap showing the four types of responses by TRPV1-GCaMP6 trigeminal neurons to brush, VF pinch, thermal (47°C), and capsaicin stimuli collected sequentially in each mouse. Color scale indicates the changes in calcium imaging intensity (ΔF/F).

TRPV1 channels are nonselective cation receptors that are gated by a broad array of noxious ligands. They act as polymodal receptors that can be independently activated by thermal, chemical, and mechanical stimulation (Cui et al., 2016). While live-imaging the TG, we applied different stimuli in the standardized sequence of mechanical, thermal, and chemical stimulation to the vibrissal pad to characterize the calcium-signaling responses of responding nociceptor populations. As expected, our results showed that TRPV1-GCaMP6f neurons respond to multiple forms of orofacial stimulation, including mechanical (brush, von Frey hair (VF)), thermal (47°C) and chemical (capsaicin) stimuli. Each of these different modes of stimulation elicited a distinct response pattern (Fig. 1C).

The application of non-noxious or noxious stimulation to the mouse vibrissal pad resulted in the robust activation of discrete populations of neurons in the mouse TG. Although some neurons displayed a response to only one type of stimulation that indicated specificity, other populations of neurons exhibited polymodal sensitivity with nociceptive responses to two or more stimuli. There were different combinations of response or non-response by individual neurons. For example, some neurons responded only to von Frey hair stimulation, others only to thermal, or only to capsaicin, yet some neurons can be activated by brush as well as von Frey, or by thermal and capsaicin stimuli (Fig. 1D).

### Increased neuronal activities in Tgp35 mice in response to painful orofacial stimuli

We next compared the response of TRPV1-linage neurons to different types of stimulation in wildtype mice compared to Tgp35 mice that have elevated Cdk5 activity due to overexpression of p35 (Harada et al., 2001; Utreras et al., 2012). We observed that the different stimuli activated various ensembles of neurons as well as inducing different calcium signal encoding (Fig. 2 A-D). Both brush and VF stimuli evoked a sharp calcium intensity peak (Fig.2 A, B), while both thermal and capsaicin stimuli resulted in a more prolonged response time (Fig. 2C, D). To quantify the differences between mice, we quantified the number of activated neurons for each stimulus and calculated the area under the curve (AUC), which combines total fluorescence intensity with temporal response time. For brush stimulation, there were no significant differences in AUC measurements between Tgp35 and control mice; however, the number of activated neurons increased significantly (Fig. 2A). In contrast, in Tgp35 mice, VF stimuli produced larger AUC Ca^2+^ intensity values while the activated neuronal population did not change (Fig. 2B). With both 47°C thermal and capsaicin stimulation, which are above the pain threshold, applied to the mouse orofacial region, total calcium intensities increased significantly in Tgp35 mice, along with enhanced numbers of activated neurons compared to controls (Fig. 2C, D). Together, these data indicate that Tgp35 mice with enhanced Cdk5 activity demonstrate increased pain stimulation induced total Ca^2+^ intensity, as well as by the number of activated neurons. In addition, differences between these measurements are stimulus dependent.

**Fig. 2.**
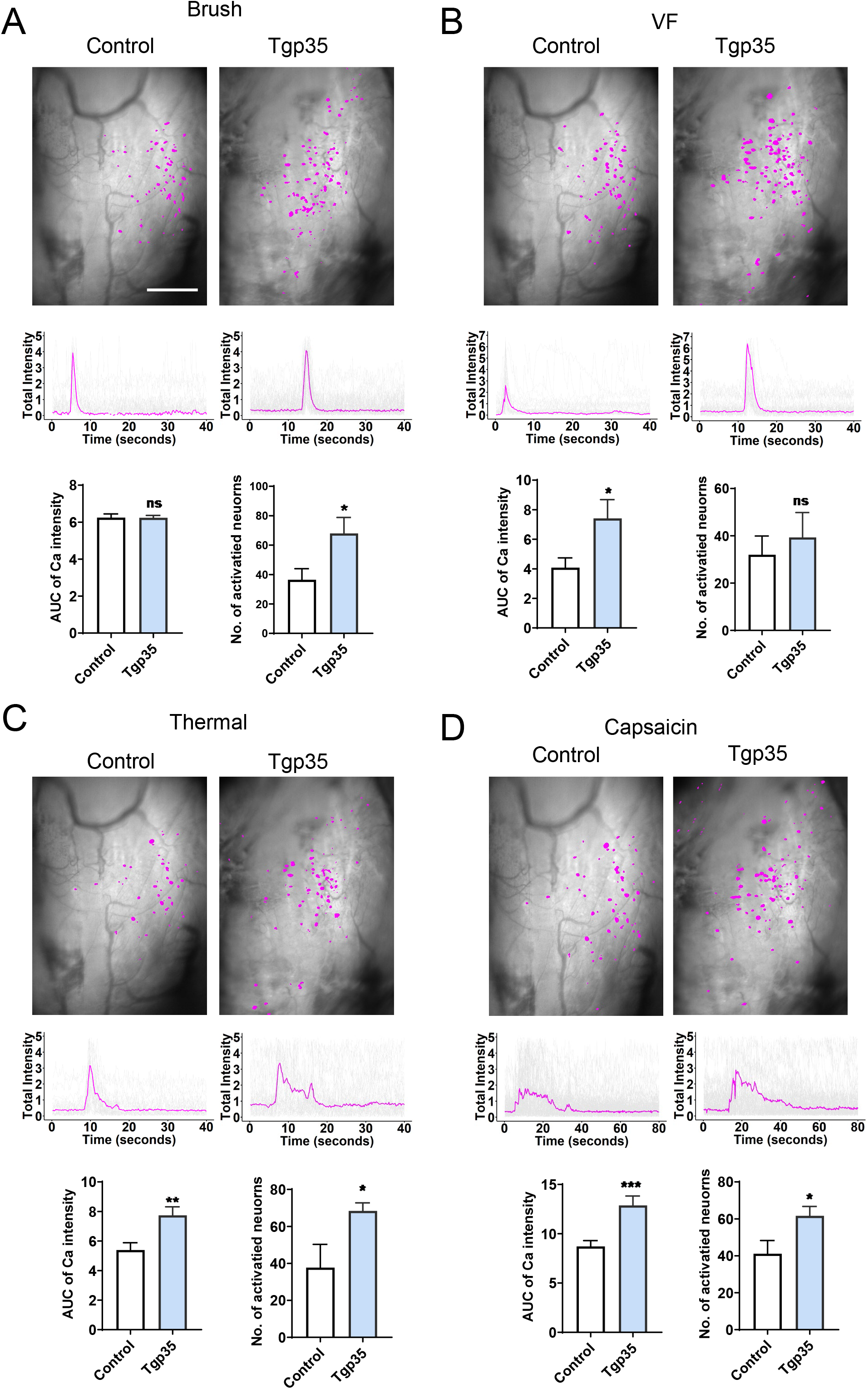
Trigeminal ganglion sensory neuronal responses to orofacial brush, von Frey hair, thermal, or capsaicin stimulation in Tgp35 mice. A. Top panels: representative imaging fields of trigeminal ganglion TRPV1-GCaMP6-expressing neurons in response to brush or von Frey hair stimulation in control and Tgp35 mice, respectively. Middle panels: representative calcium traces in response to brush stimulation in individual mice, representing typical response profiles for this stimulation. Gray lines are traces from each trigeminal ganglion neuron; magenta line shows the mean value for these traces. Left bottom panel: comparison of AUCs of calcium traces from control and Tgp35 mice in response to brush, showing no significant difference in responses to this stimulation. Right bottom panel: numbers of neurons activated by this stimulation, showing significantly more neurons responding in Tgp35 compared to control mice (n= 6). AUC, area under the curve; the data are presented as mean ± SEM, *p<0.05, ***p<0.001, scale bar, 500 μm. B. For von Frey hair (VF) stimulation (2g), the AUCs of total calcium intensity increase significantly in Tgp35 mice. There is no significant difference in numbers of neurons responding between Tgp35 and wildtype control mice. C and D. For thermal (47°C), and capsaicin stimulation (1 μM, subcutaneous injection), both AUCs and numbers of activated neurons dramatically increase in Tgp35 mice compared to wildtype mice.

### Decreased neuronal activities in p35 knockout mice in response to orofacial pain

Because of the substantial changes in neuronal firing/activity associated with Tgp35 mice, we next examined the neuronal responses of TRPV1-lineage neurons in p35 knockout (KO) mice, which have previously established diminished Cdk5 activity, in response to the same stimulations. For brush stimulation, there were no statistically significant differences in either total calcium intensity or activated neuron number between p35 knockout and control mice for this non-painful stimulus (Fig. 3A). However, in response to VF hair stimulation, AUC Ca^2+^ intensities were significantly decreased in p35 knockout mice, but the numbers of responding neurons remained similar (Fig. 3B). Both thermal and capsaicin stimulations applied to the mouse orofacial region led to decreased AUC Ca^2+^ intensities, as well as a decrease in the total number of activated neurons (Fig. 3C-D). These data, together with the findings from our Tgp35 mice, indicate a crucial role for the Cdk5-p35 pathway in regulating TG function in its response to noxious stimuli.

**Fig. 3.**
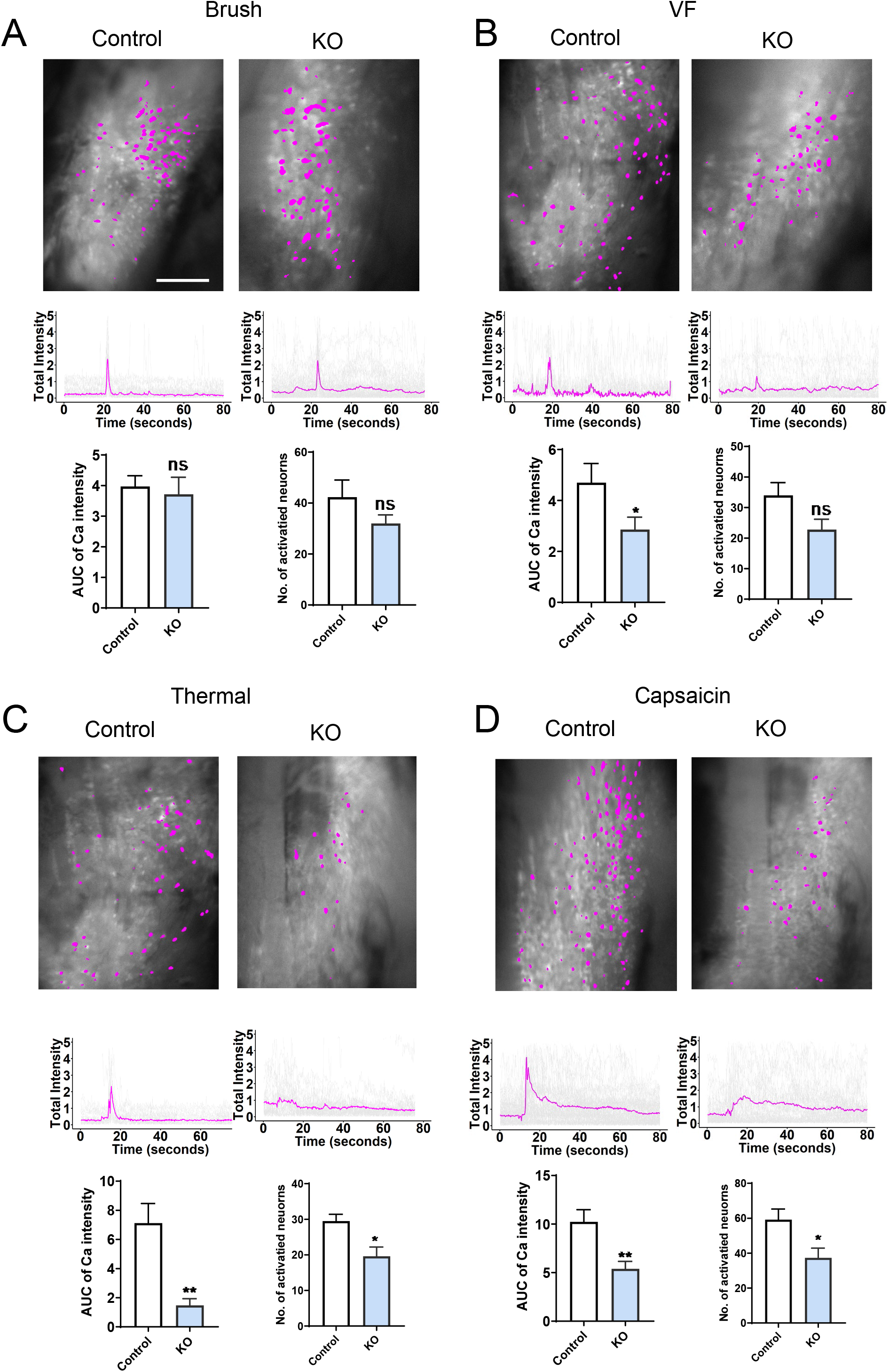
Trigeminal ganglion sensory neuronal responses to orofacial brush, von Frey hair, thermal, or capsaicin stimulation in control vs. p35 knockout (KO) mice. A. Top panels: representative imaging fields of trigeminal ganglion Trpv1-GCaMP6f expressing neurons responding to brush stimulation in control versus p35 knockout mice. Middle panels: representative calcium traces in response to brush stimulation in individual mice. Gray lines are traces from each trigeminal ganglion neuron; magenta line is the mean value for these traces. Left bottom panel: comparison of AUCs (areas under the curve) of calcium traces from control and p35 mice in response to brush, showing no significant difference in responses to this stimulation. Right bottom panel: numbers of neurons activated by this stimulation, showing decreased trend but no significant difference in neurons responding in p35 knockout compared to control mice (n= 6). the data are presented as mean ± SEM, *p<0.05, ***p<0.001, scale bar, 500 μm. B. For von Frey hair (VF) stimulation (2g), the AUCs of total calcium intensity decrease significantly in p35 knockout mice, but with no significant difference in numbers of neurons responding. C and D, For thermal (47°C) and capsaicin stimulation (1 μM, subcutaneous injection), both AUCs and numbers of activated neurons dramatically decrease in p35 knockout mice compared to wildtype mice.

**Fig. 4.**
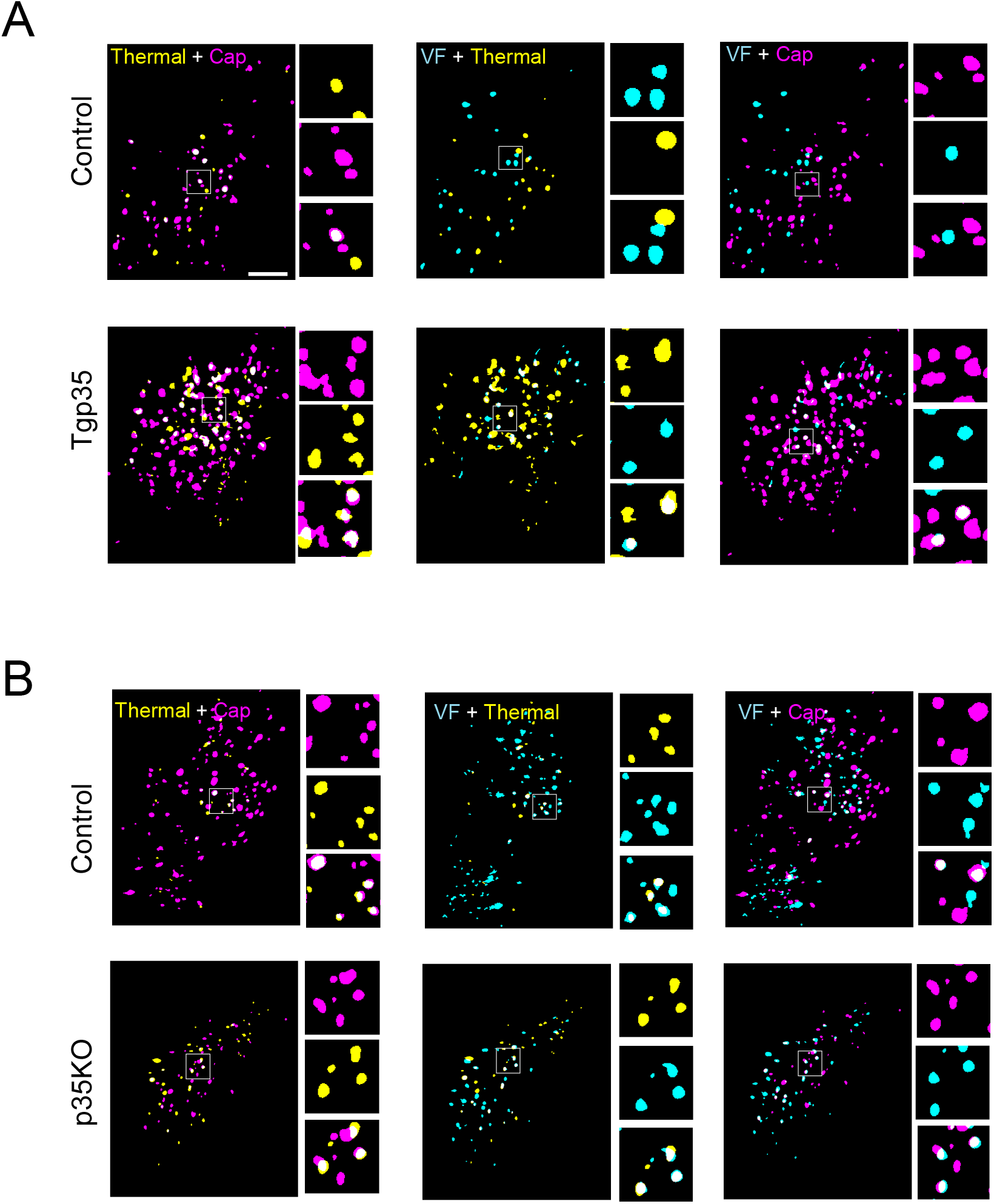
Representation of specific and polymodal neuron activation in Tgp35 and p35 knockout mice. A. Left panel: Color-coded map of a representative field showing distribution of neurons activated by either capsaicin (magenta) or thermal (yellow) or by both stimulations (white) in control and Tgp35 mice. The insets show magnified views of white-boxed region. Middle and right panels show color-coded maps of the distributions of neurons responding to von Frey hair (VF, cyan) and thermal (yellow) or capsaicin (red) stimulation, respectively; white indicates overlap. Scale bar, 500 μm. B. Left panel: Color-coded map of neurons activated by either capsaicin (magenta) or thermal (yellow), or by both stimulations (white) in control vs. p35 knockout mice. On the right side is zoom in version of squared area, illustrates thermal, capsaicin stimulation and the merge of two stimulations. Middle and right panel is color-coded map showing distribution of neurons respond to VF and thermal, VF and capsaicin stimulations respectively.

### Specific versus polymodal TRPV1-linage nociceptors in Tgp35 and P35 knockout mice

We next characterized the specific (firing in response to only a single stimulus) and polymodal neurons in TRPV1-GCamp6f mice in response to noxious stimuli. Fig. 4A and B illustrates these findings visually, showing color-coded merged images for the same microscope imaging field with various activated neurons responding to only single, two, or three stimuli in Tgp35 and p35 knockout mice. Venn diagrams (Fig. 5A) for control and Tgp35 mice revealed alterations in the proportion of neurons that demonstrate a specific or a polymodal (activation by 2 or more stimuli) response. In the Tgp35 mice with elevated Cdk5 activity, there were notably substantial increases in polymodal neurons in response to all noxious stimuli: 206%, 55%, and 240% increases in VF, thermal, and capsaicin stimuli, respectively (Fig. 5B Chi-square test, *p<0.05). Enhanced Cdk5 activity also significantly increased the proportion of specific neuronal nociceptor response to thermal stimulation (Fig. 5C). In terms of total specific/unimodal nociceptors in Tgp35 mice, there was a significant decrease, while the percentage of polymodal nociceptors correspondingly increased (Fig. 5D). These data provide evidence that increased Cdk5 activity results in cross-excitation of specific nociceptors, which in turn increases the number of polymodal nociceptors. In p35 knockout mice, however, the ratio of specific/unimodal to polymodal nociceptors remained same, i.e., there was no switch to increased specificity in responses to any specific mode of pain stimulation (Fig 5E-H). Decreased Cdk5 activity only affected integrated Ca^2+^ intensity but had no effect on the activation of specific nociceptors and polymodal nociceptors.

**Fig. 5.**
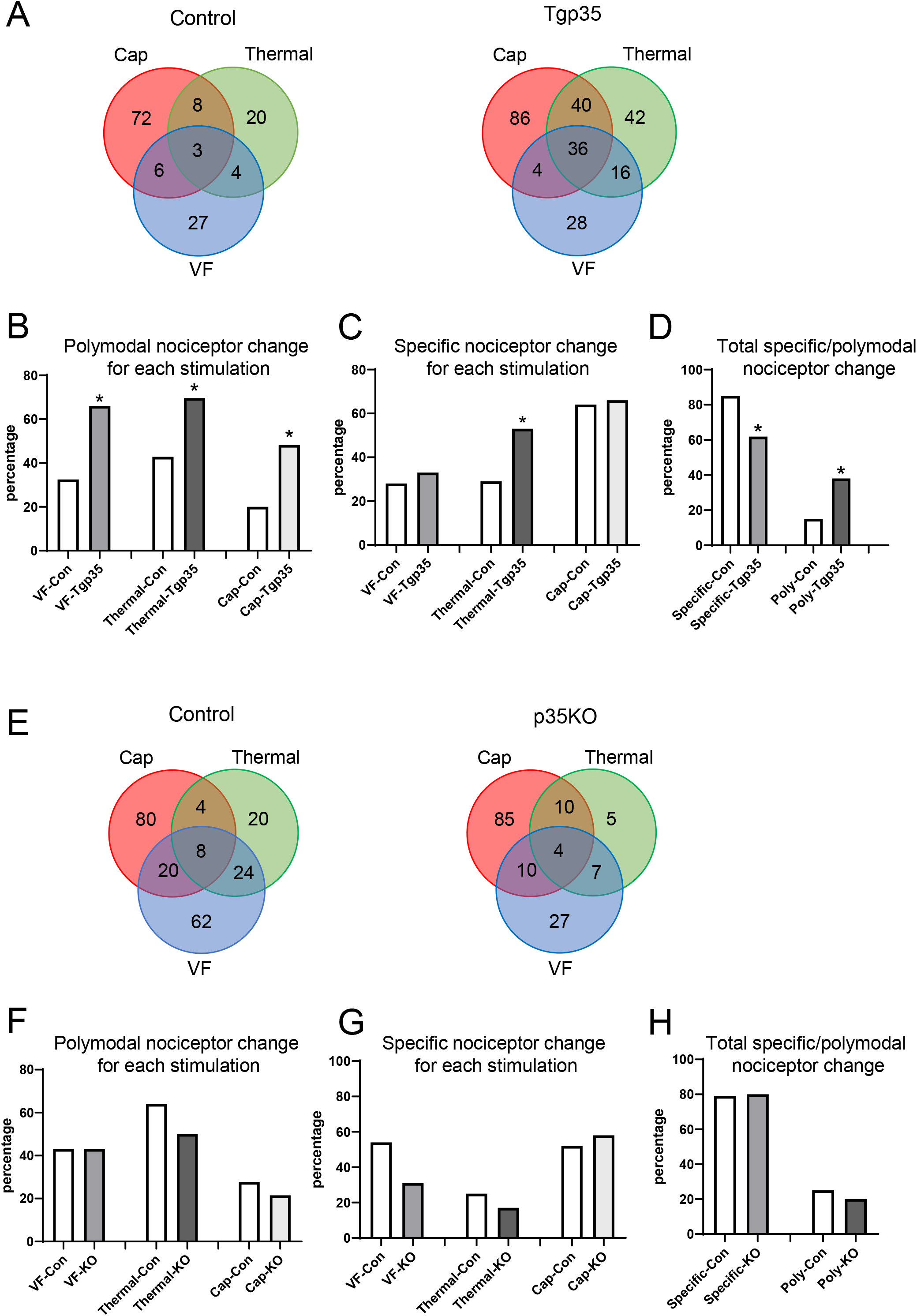
Altered distribution of polymodal sensory neuron signaling in trigeminal ganglia of Tgp35 and p35 knockout mice. A. Venn diagrams illustrating the number of specific versus polymodal (overlapping circles) TRPV1-linage trigeminal ganglion neurons responding to von Frey hair (VF, blue), thermal (green), and capsaicin (red) stimulation. n=3 mice, 392 neurons. B. The percentage change of polymodal nociceptor for VF, thermal and capsaicin stimulation. There is a significant increase in polymodal nociceptor for each stimulation. C. Percentage change of specific nociceptor neurons that respond to VF, thermal, or capsaicin. There is a significant increase in neuronal response to thermal stimulation compared to control mice by chi-square test. D. Altered percentages of specific versus polymodal (respond to 2-3 stimuli) nociceptor neurons responding to sequential von Frey mechanical, thermal, and capsaicin stimulation in control versus Tgp35 mice. E. Venn diagrams illustrating the numbers of specific and polymodal of TRPV1-linage trigeminal ganglion neurons responding to von Frey mechanical (blue), thermal (green), or capsaicin (red) stimulation in control vs. p35 knockout mice. n=3 mice, 366 neurons. F. The percentage change of polymodal nociceptor for each stimulation. G. The percentage distribution of specific nociceptor neurons responding to von Frey, thermal, or capsaicin stimulation. H. Summary of the percentages of specific and polymodal nociceptors responding to von Frey, thermal, or capsaicin stimulation, indicating no significant change in the proportions of signaling from specific compared to polymodal nociceptor neurons.

### Effects of TFP5 on complete Freund’s adjuvant (CFA)-induced inflammatory pain

After establishing that Cdk5 activity affects trigeminal ganglion peripheral neuronal signaling, we tested whether a Cdk5 inhibitor can suppress inflammatory pain signaling at a peripheral site, rather than at the CNS level. We used the classical orofacial CFA mouse model for animal modelling of inflammatory disorders – an accepted model of orofacial inflammatory hyperalgesia and allodynia for nearly five decades (Liao et al., 2017; Morgan and Gebhart, 2008). Each mouse first received an injection of 50 μl CFA into the vibrissal pad. After 6 hours, live imaging was conducted to monitor neuronal responses after TFP5 peptide (80 mg/kg) was applied directly to the trigeminal ganglion. We observed that in control CFA-induced inflammation mice, there is a polymodal neuronal response to both non-noxious (brush) and noxious (capsaicin) stimulation that represents inflammation-induced hyperalgesia and allodynia. Fig. 6A illustrates examples of color-coded activated neurons responding to capsaicin or brush stimulation with scrambled (Scb) control and TFP5 peptide, respectively, in the CFA-induced inflammation mice. The merged image shows polymodal neurons (yellow) responding to both brush and capsaicin stimulation. Additionally, the amplitude of neuronal response to both brush and capsaicin decreased significantly after TFP5 treatment compared with treatment with the scrambled peptide control; moreover, AUC Ca^2+^ intensities also showed significant reductions after TFP5 application (Fig. 6B). It is noteworthy that the number of neurons responding to both brush and capsaicin also decreased in TFP5-treated mice. We compared the percentage of this class of polymodal neurons in control mice, CFA-induced inflammation mice, and the TFP5 treatment group. There was a dramatic increase in the proportion of polymodal neurons after CFA injection compared with control, and in the presence of TFP5, it was reduced substantially (Fig. 6C). Together, these data suggest that Cdk5 inhibitor can downregulate the number of nociceptors responding to both brush and capsaicin and decrease the integrated Ca^2+^ transients in response to both stimulations in inflamed mice. Thus, Cdk5 inhibitor can reverse the allodynia induced by inflammation.

**Fig. 6.**
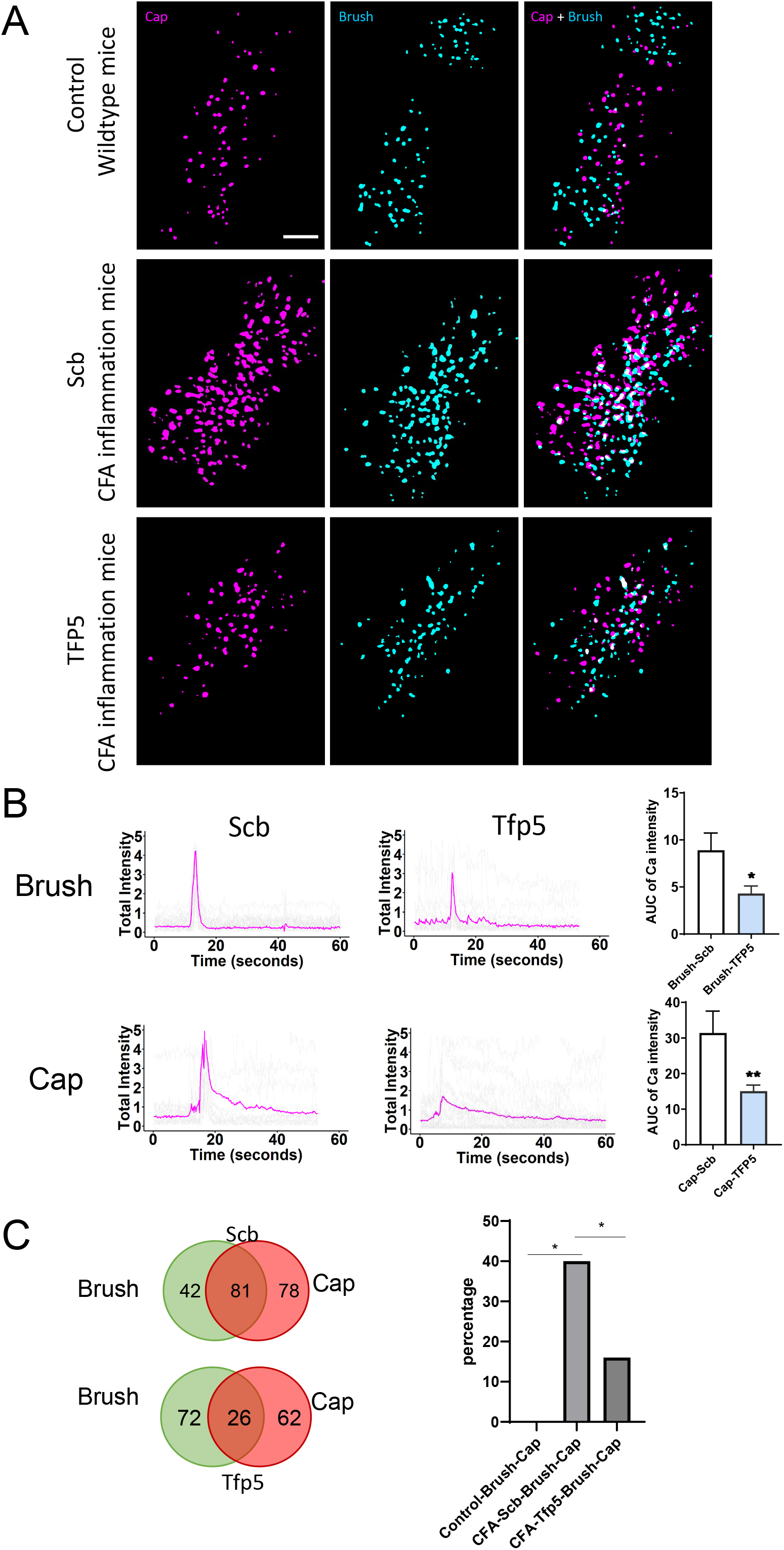
Effects of direct application of the Cdk5 inhibitor TFP5 to trigeminal ganglia on neuronal responses to orofacial brush and capsaicin in mice with orofacial inflammation. A. Representative color-coded map illustrating neuronal responses to capsaicin (magenta) and brush (cyan) stimulation in scrambled (Scb) control peptide and TFP5 treated mice after CFA injection to induce inflammation. The right panel shows the merge of two stimulations, where white indicates overlap. B. Left panels show representative calcium traces responding to brush or capsaicin stimulation treated with scb control or TFP5 peptides. Right panels show graphical summaries comparing AUCs for both stimulations in Scb and TFP5 mice with a substantial decrease after TFP5 treatment compared to the Scb (scrambled) peptide control. C. Venn diagrams showing the distribution of specific and polymodal neuronal responses to brush and capsaicin stimulation. n=3 mice, 361 neurons. D. Altered percentage of polymodal nociceptors responding to both brush and capsaicin stimulation. There is a robust increase of polymodal nociceptors after CFA-induced inflammation in the orofacial region of mice, and a significant decrease after TFP5 treatment.

## Discussion

In this study, we used intravital imaging in mice to compare the regulation of trigeminal ganglion (TG) neuronal signaling in responses to four different stimuli. Key findings included the following: Cdk5 activity regulates both the intensity of Ca^2+^ signaling and its specificity, (i.e., whether the neuronal signaling is specific or polymodal). Inflammation could significantly enhance TG polymodal signaling, and a peptide inhibitor of Cdk5 was able to suppress both the intensity and extent of polymodal signaling associated with inflammatory pain and experimental allodynia when administered either peripherally to the TG or intraperitoneally.

Although facial and oral pain affects millions of patients, the mechanisms underlying this pain sensed by the TG are still incompletely understood. Because TRPV1 receptors are highly expressed in TG nociceptors, we focused our attention on these cells as possible targets for therapeutic intervention using direct assays of neuronal calcium signaling by in vivo live GCaMP6f imaging.

We characterized new patterns of specific and polymodal signaling in the trigeminal ganglia of mice by direct intravital imaging of the dynamics of calcium signaling in response to the following four different stimuli in mice. Mechanical facial stimuli included light touch (brush) and pinch via calibrated point pressure using von Frey filaments, whereas painful thermal stimulation was by direct contact to the vibrissal pad, and chemical stimulation of pain was evaluated using capsaicin. Different stimulations induced distinct neuronal integrated calcium transients, which were generally consistent with previous studies (Ghitani et al., 2017; Leijon et al., 2019).

We directly compared the regulation of these modes of signaling in TRPV1-linage neurons after modulating activity levels of the Cdk5. Our previous behavioral study had shown that mice with increased Cdk5 activity show lower tolerance to TRPV1-mediated painful stimuli, whereas mice with reduced Cdk5 activity have higher tolerance to the same stimulus (Jendryke et al., 2016; Prochazkova et al., 2013). However, whether Cdk5 directly affected primary sensory neurons or acted on the CNS remained elusive. In this study, we established that Cdk5 can regulate both the amplitude of pain signaling and the number of responding neurons. In contrast, altering Cdk5 activity had no apparent effect on the response to non-noxious mechanical stimulation (brush), except for some increased numbers of responding neurons after Cdk5 hyper-activation. Interestingly, we were able to demonstrate altered ratios of specific and polymodal responses to all stimuli -- more neurons began to exhibit signaling to more than one stimulus after elevation of Cdk5 activity. Increased Cdk5 induced cross-excitation of primary sensory neurons in the trigeminal ganglion with enhanced signaling due to light touch (brush), indicating hyperalgesia and allodynia. Conversely, genetic ablation of Cdk5 resulted in fewer responding neurons and more unimodal signaling when sensing thermal and noxious mechanical pain (pinch).

In response to facial inflammation, afferent nociceptors can be stimulated by activity in low-threshold mechanoreceptors, resulting in cross-excitation between A and C fibers in sensory neurons to induce allodynia (Amir and Devor, 2000). TRPV1-positive neurons are essential for the development of mechanical allodynia. In rats exhibiting neuropathic pain, the TRPV1-positive neurons mediate the most sensitive aspects of mechanical allodynia (Tender et al., 2008). In line with these studies, our live imaging illustrated that in our mouse model of facial inflammation, both separate and overlapping-modality Trpv1^lin^ TG neurons were activated in response to noxious or previously non-noxious stimulation, i.e., enrichment for polymodal sensory neuronal responses to both brush and capsaicin.

Compared with control mice, the percentage of polymodal nociceptor cell responses to brush and capsaicin stimuli increased dramatically from 0.3% to 39% (Fig. 6D). This finding represents the first demonstration to our knowledge that at the level of TG calcium neuronal signaling during the inflammatory process, responses to noxious stimuli are enhanced or pain is triggered by normally innocuous stimuli. (Smith-Edwards et al., 2016) have demonstrated inflammatory pain could shift the balance of sensory neuronal signaling in DRG, which is consistent with our findings. TRPV1 acts as a key receptor in nociceptive neurons, and its function is strongly affected by Cdk5 phosphorylation that can lead to hyperalgesia and allodynia (Jendryke et al., 2016; Simonic-Kocijan et al., 2013). Cdk5 regulates TRPV1 membrane trafficking in an inflammatory model to increase sensitivity to pain (Xing et al., 2012). It has also been shown that intrathecal administration of a Cdk5 inhibitor alleviates inflammatory heat hyperalgesia in rats (Liu et al., 2015).

Here, we provide in vivo evidence that a Cdk5 inhibitor peptide TFP5 can act directly on peripheral nerve pain signaling at the trigeminal ganglion. Direct treatment of trigeminal ganglia with TFP5 not only downregulated the polymodal nociceptor response to brush and capsaicin, but also decreased the integrated calcium signaling for both stimuli. Virtually identical results were obtained by intraperitoneal injection of this inhibitor. By suppressing pain signaling in peripheral TG neurons, inhibition of Cdk5 could provide an approach to suppressing pain at a peripheral level rather than at the usual central (CNS) level.

Our current study directly demonstrates the effect of Cdk5 activity on facial pain signaling in primary sensory neurons, suggesting a new approach or pathway to alternative pain therapeutics. Targeting only peripheral TG neurons could produce significant analgesic effects. This finding might ultimately have substantial translational impact, since most of current pain medications are not selective for primary neurons and affect their targets in multiple tissues particularly the central nervous system, often leading to serious side effects. Targeting Cdk5 activity at TG primary sensory neurons could offer better and safer treatments for orofacial pain.

## Acknowledgments

We thank the National Institute of Dental and Craniofacial Research (NIDCR) Imaging Core for imaging technical support and the NIDCR Veterinary Resources Core for animal support. We thank Dr. Alexander T. Chesler, Dr. Nima Ghitani and Bradford Hall for their help. We thank Dr. Shaohe Wang for his input for machine learning scripts. This research was supported by the Intramural Research Program of the NIH, NIDCR.

## Authors Contribution

Conceptualization, M.H., A.D.D. A.B.K. and K.M.Y.; methodology, M.H., A.D.D. A.B.K. and K.M.Y.; investigation, M.H., A.D.D. A.B.K. and K.M.Y.; writing—original draft, M.H., A.D.D.; writing—review and editing, K.M.Y., M.H., A.D.D. A.B.K. and K.M.Y.; funding acquisition, A.B.K., K.M.Y

## STAR Methods

### Animals

Mice were housed in a temperature- and light- controlled room (23±2°C and 12 h light/dark cycle, respectively) with ad libitum food (2918 Teklad global 18% protein rodent diet, Envigo) and water. TRPV1-Cre (stock number 017769) and GCaMP6f (stock number 028865) strains were purchased from Jackson Laboratories. Mice with p35 knockout or transgenic overexpression, i.e., p35^-/-^ background and tgp35 mice, were generated in our laboratory(Pareek and Kulkarni, 2006). P35^-/-^ mice were maintained in a C57BL6/129SVJ background. Tgp35 mice and wild-type littermate controls were maintained in an FVBN background. For imaging, mice were crossed to generate a GCaMP6 and TRPV1-Cre line, which was then maintained with p35^-/-^ or tgp35 backgrounds. For all experiments, age-matched wild-type littermates served as controls. All experimental procedures were approved by the Animal Care and Use Committee of the National Institute of Dental and Craniofacial Research, National Institutes of Health.

### Surgery

Surgical procedures were conducted as described previously (Ghitani et al., 2017; Hu, 2019). In brief, mice at 8-10 weeks of age were anesthetized with inhalational Isoflurane (2%)/oxygen mix and positioned in a custom-built stereotactic frame with heads immobilized using ear bars. Continuous flow of isoflurane/oxygen was provided through a nosecone (Braintree Scientific, Inc.). Body temperature was maintained at 37°C ± 0.5°C using a heat mat (CWE, Inc.), and temperature was monitored by a mouse rectal temperature probe throughout surgery and imaging. Under a dissecting stereoscope (Leica), scissors (Fine Science Tools) were used to remove skin from the top of the head, then connective tissue was removed to expose the skull, followed by partial decerebration surgery to obtain optical access to the trigeminal ganglion (Hu 2019). The cranium was repeatedly washed and bathed in HEPES buffer (160 mM NaCl, 6 mM KCl, 13 mM glucose, 13 mM HEPES, 2.5 mM CaCl_2_, 2.5 mM MgCl_2_ with pH adjusted to 7.2 using 10N NaOH). A rubber O-ring (9 mm diameter, RT Dygert) was glued over the opening in the skull using a cyanoacrylate-based adhesive. A custom-made stabilization bar securely mounted to the frame was attached to the O-ring and skull using dental cement.

### Inflammation model

Mice were anesthetized with 2% isoflurane mixed with oxygen and were injected subcutaneously with 15 μl of complete Freund’s adjuvant (CFA) solution (Sigma-Aldrich), or saline into the lateral facial skin using a 31-gauge needle (BD Ultra-Fine 6 mm needle). 6 hours after CFA or saline injection, mice were anesthetized and transferred to the imaging stage.

### Calcium imaging data collection and analysis

After surgery, the mouse was transferred to the stage of an epifluorescence inverted microscope (IX-71, Olympus) equipped with a 4X, 0.28 NA air objective attached to a inverting device (LSM Technologies). Illumination was provided by a 130 W halogen light source (Olympus), using a standard GFP excitation/emission filter cube. Imaging was performed using an Orca Flash 4.0 CMOS camera (2016 Model, Hamamatsu), imaging at a 5 Hz frame rate, using Micro-Manager acquisition software to collect data.

Imaging data were analyzed using custom macro algorithms in ImageJ (NIH). For each field of view, all image stacks were first aligned for motion correction (MetaMorph, Molecular Devices). To identify neurons, maps of peak activity (maximal pixel intensity over mean pixel intensity) was median filtered, thresholded and separated by watershed segmentation to create regions of interest representing active neurons. A 2-pixel width ring around the circle were considered as local background. Fluorescence traces for each region of interest were subtracted from local background intensity.

### Thermal, light touch, von Frey hair, and capsaicin stimulations

All stimuli were applied while the mouse was anaesthetized and under continuous live imaging. For thermal stimulation, we use a copper tubing loop stimulator connected to a circulating water bath (LW Scientific, Italy). The temperature was maintained at 47.0°C (±0.5°C) to reach the hot nociceptive threshold. The stimulator was applied to the center of the right vibrissal pad of the mouse. Mechanical stimulation included both brush (light touch) and von Frey hair stimulation. Brush was performed using a cotton swab to gently brush the mouse right vibrissa pad. Von Frey hairs consisted of plastic monofilaments of equal length and varying diameters for which the force required to bend each filament was calibrated (Vos et al., 1994), We used 2g force von Frey filaments applied to the center of the right vibrissal pad.

We also examined the effects of low (1 μM) and high (10 μM) doses of capsaicin on TG neuronal activity. Capsaicin (Millipore Sigma) was injected subcutaneously through a 31-gauge needle (BD Ultra-Fine 6mm needle) into the center of the right vibrissal pad.

### Statistical analysis

All data are expressed as mean ± SEM, and n represents the number of mice analyzed. The statistical evaluation was performed with GraphPad Prism 8 software (GraphPad, San Diego, CA). Statistical differences between the groups were assessed using a two-tailed unpaired *t* test with Welch’s correction. A statistically significant difference was defined as p < 0.05. Error bars indicate SEM.

